# Recent lifestyle change impacts sleep and circadian rhythms among the Indigenous peoples of Peninsular Malaysia

**DOI:** 10.1101/2025.07.26.666903

**Authors:** Kathleen D. Reinhardt, Thomas S. Kraft, Amanda J. Lea, Ian J. Wallace, Yvonne A.L. Lim, Colin Nicholas, Tan Bee Ting A/P Tan Boon Huat, Kar Lye Tam, Steven K.W. Chow, Izandis bin Mohd Sayed, Kamal Solhaimi Fadzil, Michael C. Antle, Vivek V. Venkataraman

## Abstract

Sleep disorders are rising globally, but their lifestyle causes remain unclear. We recorded sleep-wake patterns via actigraphy from 1036 Orang Asli adults across 12 communities in Peninsular Malaysia undergoing market integration, marked by changes in permanent infrastructure (electricity and housing), digital technologies (smartphones), and labor practices (i.e., wage labor). We evaluated associations with sleep timing (onset, offset and regularity), quality (nighttime awakenings and waking after sleep onset) and quantity (sleep duration), while accounting for age and sex. Delayed and destabilized sleep timing was observed in communities with powerline access, also resulting in shorter sleep duration; paradoxically, it also improved sleep quality, suggesting increased homeostatic pressure. Age and sex were strong and consistent predictors of sleep variation: older adults had earlier, shorter, and more consistent, consolidated sleep patterns. Men displayed later and shorter sleep patterns than women, likely reflecting gendered divisions of labor among the Orang Asli. Despite averaging relatively few hours slept (6 hrs), Orang Asli exhibited relatively efficient sleep, potentially challenging the notion that longer sleep is universally beneficial. These findings underscore the complex interplay of biology, ecology, and culture in shaping sleep and circadian rhythms.

**Significance:** A comprehensive cross-sectional study of sleep across a pronounced lifestyle gradient among Malaysia’s Indigenous Orang Asli populations reveals new insights into the drivers of human sleep and circadian rhythms. Lifestyle changes with market integration, particularly access to electricity, resulted in delayed bedtime and shortened sleep duration, yet enhanced sleep consolidation. Consistent with cross-cultural evidence, aging resulted in earlier bedtimes, earlier rising times, and less sleep. Our findings contribute to debates about the adaptability of human circadian rhythms and challenge universal models of optimal sleep duration.

## INTRODUCTION

Chronic sleep disorders are a pervasive public health concern in industrial and post-industrial societies, with robust epidemiological evidence linking them to a heightened risk of major non-communicable diseases (Hirshkowitz et al., 2015; Hale et al., 2020; Zheng et al., 2024). These disorders often present as reduced total sleep duration (impaired quantity), diminished sleep efficiency (compromised quality), delayed sleep phase (misaligned timing), or a combination of these disturbances (Chattu et al., 2018; Gradisar et al., 2013; Knutson et al., 2017). While the high prevalence of chronic sleep disorders in industrial and post-industrial societies is widely hypothesized to stem from recent lifestyle changes, the specific determinants remain incompletely understood. However, there are three particularly conspicuous candidates, all of which have the potential to disrupt sleep homeostasis primarily by dysregulating the endogenous circadian clock. These include features of modern technology, the built environment, and labor practices.

The endogenous circadian clock, which governs near-24-hour cycles of physiological and behavioral processes, relies on external zeitgebers (circadian cues)—most critically, the light-dark photoperiod—to maintain synchrony with the solar day (Pittendrigh, 1981; Roenneberg et al., 2007, Roenneberg et al., 2013). Yet many modern technologies, including electric lighting, digital screens, and round-the-clock connectivity, prolong exposure to artificial light at night, increasing the risk of circadian misalignment (Chang et al., 2015; Gronfier et al., 2007). In particular, electric lighting extends the perceived day as well as facilitates pre-dawn activity, potentially delaying sleep onset and leading to earlier sleep offset (Peirson et al., 2018). Compounding this disruption, the built environment of industrial and post-industrial societies often diminishes natural zeitgebers: architectural designs limit natural daylight exposure, climate control minimizes temperature fluctuations, and sound insulation muffles ambient noise, all of which may impair circadian entrainment (Mistlberger & Skene, 2005; Roenneberg & Merrow, 2007; Wright et al., 2013). Further exacerbating this issue, contemporary labor patterns—marked by shift work, long hours, awkward schedules, and repetitive tasks—frequently oppose innate sleep-wake rhythms (Fischer et al. 2021; McHill et al., 2014; Vetter et al., 2018).

Nevertheless, the hypothesis that chronic sleep disorders are driven by recent shifts in technology, the built environment, and occupational demands is not unequivocally supported by comparative studies of sleep among non-industrial societies, which document sleep patterns resembling those of industrial and post-industrial societies in some key respects. For instance, research among hunter-gatherers in Africa and forager-horticulturalists in South America revealed average sleep durations comparable to those observed in the United States (de la Iglesia et al., 2015; Pilz et al., 2018). A potential explanation for this paradox is that modern lifestyle factors exert heterogeneous effects on sleep, with some influences being negative, neutral, and/or even beneficial. For example, features of modern housing—such as attenuated exposure to external zeitgebers (e.g., ambient light, temperature variability)—potentially mitigate premature sleep offset, thereby extending sleep duration while also mitigating nighttime disruptions during sleep. A comprehensive investigation into the concurrent effects of technology, the built environment, and labor practices on sleep timing, quality and quantity in a traditionally non-industrial society undergoing lifestyle change could help elucidate the complex trade-offs imposed by modern lifestyles. However, to date, no such study has been conducted.

Here, to better understand the impact of modern lifestyle factors on sleep variation, we investigate determinants of sleep patterns among the Orang Asli, the Indigenous peoples of Peninsular Malaysia (Endicott, 2016). Traditionally rainforest-dwelling hunter-gatherers and subsistence farmers, the Orang Asli have experienced varying degrees of lifestyle change in recent decades due to acculturation, ecological degradation, and integration into the market economy (Wallace et al., 2022; Watowich et al., 2024). This transition has produced a broad spectrum of lifestyles, ranging from communities maintaining traditional bamboo dwellings with minimal electricity or market engagement to urbanized individuals residing in concrete houses, engaged in wage labor, and with easy access to modern technologies. Using a large sample of Orang Asli spanning this lifestyle gradient and leveraging actigraphy measurements of sleep patterns, we test the overarching hypothesis that recent lifestyle shifts—specifically access to modern technologies, changes in the built environment, and engagement in wage labor—impact sleep timing, quality, and quantity. Our general prediction is that modern lifestyle conditions negatively affect sleep patterns, though we recognize that some positive effects are possible (Samson & McKinnon, 2025). Beyond elucidating the effects of modern lifestyle factors on sleep patterns, this study contributes to growing evidence of sleep among non-industrial societies— whose lifestyles resemble those of our ancient evolutionary ancestors—thereby offering critical insights into the evolution of human sleep.

## METHODS AND MATERIALS

### Overview of study design

This study was conducted as part of the Orang Asli Health and Lifeways Project (OA HeLP), an interdisciplinary team of anthropologists, physicians, biomedical researchers, ecologists, and human rights activists interested in the causes and consequences of Orang Asli health (Wallace et al 2022). Since 2020, our team has worked in numerous Orang Asli communities across Peninsular Malaysia, spanning the gradient from remote rainforest camps and villages to urban and peri-urban settlements.

In each community, the OA HeLP team collects biological samples and data, conducts in-depth ethnographic interviews, and provides free medical care through team physicians. Interviews cover a wide range of topics, including: (1) their source of electricity, if any; (2) whether they own specific light-emitting devices such as smartphones and televisions; (3) the construction of their housing; and (4) the extent to which they engage in wage labor versus traditional subsistence activities. Additionally, once the research team has completed their work in a community, each study participant is given a triaxial accelerometer (Axivity AX3, Axivity Ltd, UK) to wear for up to 10 days on their non-dominant wrist to measure their sleep (and physical activity) patterns. Accelerometers are configured to record information at a sampling frequency of 100 Hz with a dynamic range of 8 *g*. A community volunteer later collected the devices and eventually returned them to the research team. Accelerometry is restricted to periods when researchers are not working in the community to ensure that the presence of our team does not alter behavior. The use of accelerometry has been extensively validated with polysomnography as a method for measuring human sleep-wake patterns in both field and laboratory settings.

Data included in the present study derive from 1107 Orang Asli aged 18–91 years from 12 communities studied between 2020 and 2024, which exhibit substantial variation in access to modern technology, built environment, and labor practices (Figure 1, Table S1). The sample includes individuals from multiple Orang Asli tribes, including the Batek, Jahai, Semai, Temiar, and Temuan.

**Figure 1.**
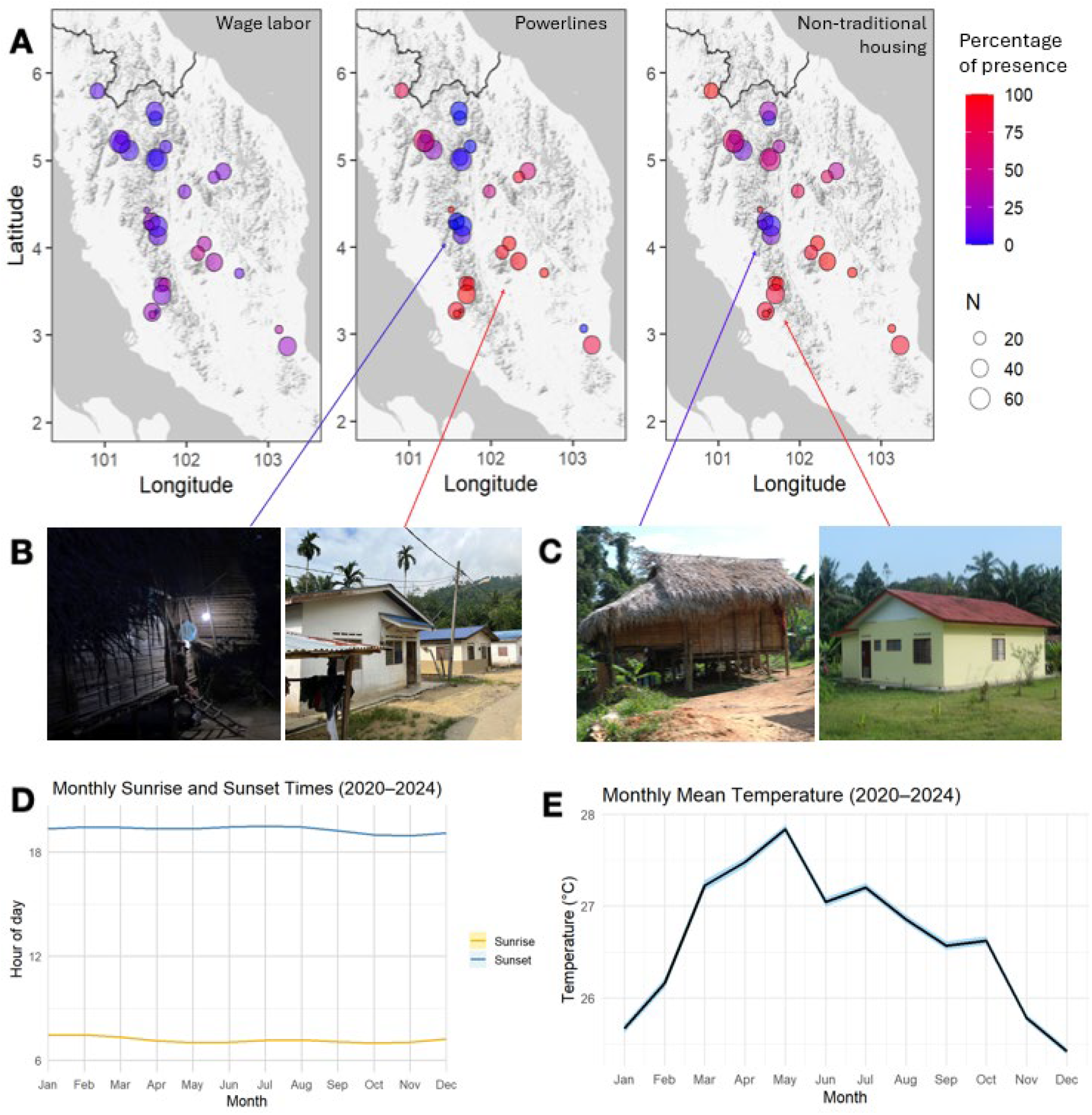
Variation in wage labor, electricity, housing type and climate across study locations. (A) Participant Orang Asli villages across Peninsular Malaysia, with the number of participants per study location corresponding to the size of the circle. Circles are shaded by the number of individuals at each location that reported participating in wage labor, having access to power lines, or living in non-traditional (i.e., wood or concrete) housing. (B) Examples of solar power electricity (left) and community powerlines (right). (C) Examples of a traditional thatch roof house (left) and a non-traditional concrete house (right). (D) Monthly average sunrise (yellow) and sunset (blue) times from 2020-2024 in Kuala Lumpur, Malaysia (Time and Date AS, retrieved July 09, 2025). (E) Monthly mean near-surface air temperature (°C) from 2020-2024 in Peninsular Malaysia (4.8° N, 102° E). The curve shows a loess-smoothed trend with 95% confidence intervals (blue ribbon), based on hourly ERA5 data at 2 m height (Hersbach et al. 2020).

### Accelerometry data processing

We extracted raw triaxial accelerometry data using OmGUI (Newcastle, UK). Accelerometry data were processed to generate summary metrics of sleep outcomes using the GGIR package (Migueles et al., 2019; van Hees et al., 2015) in R]. Sleep period time detection was performed using the Heuristic algorithm looking at Distribution of Change in Z-Angle (van Hees et al. 2018) for all days exceeding a threshold of at least 16 hours of detected wear time in the noon-noon period encompassing a given night. A full range of the GGIR parameters employed in this study is included in Supplementary Materials.

Accelerometry deployments were not accompanied by sleep diaries or direct observation; thus, all post-processed actigraphy was visually inspected (Winnebeck et al., 2018). Nights were excluded if they exhibited implausible sleep durations (<2 hours or >12 hours), >20% continuous non-wear time during the sleep period, or highly irregular sleep–wake patterns (Littner et al., 2022; van Hees et al., 2015). We also identified nights showing two distinct sleep bouts with sustained activity between them, known as biphasic sleep or “two sleeps” which can be typical among non-industrial societies (Samson et al., 2017a; Smit et al., 2019; Yetish et al., 2015). As merging these bouts can provide misleading or biased interpretations of sleep quality and architecture, we omitted all nights with biphasic sleep (n= 333).

### Exposure and outcome variables

Exposures of interest included modern technologies (electricity access, ownership of light-emitting technology), built environment (housing type), and labor practices (engagement in wage labor). For electricity access, we categorized each participant as having access to solar power, a gas-powered generator, and/or power lines connected to an electrical grid, or no access to electricity. Access to solar power typically means owning one or more small solar lamps, which have a limited operation time, thus being qualitatively similar to having no electricity source. When present, generators are used as a community resource, with lines running to some or all houses. Thus, most individuals use it to power lights in their homes. Generators are used intermittently due to constraints on fuel availability and are typically turned on shortly after sunset and turned off from 23:00 - 24:00. In communities with power lines, virtually all individuals have easy access to electricity throughout the night and day. For light-emitting technology, we focused on ownership of a smartphone. Although we also asked participants about television ownership, we chose not to test the effects of such ownership on sleep, since most communities have a small number of televisions and they are watched communally, making TV ownership an unreliable proxy for television watching. Smartphones, in contrast, are personally owned. For each participant, housing type was categorized based on construction materials as either a traditional house made from bamboo (which typically have thatch roofing) or a non-traditional house made from wood or concrete (which typically have zinc roofing). Most concrete houses have a standardized design and are built by the government in an effort to resettle Orang Asli into authorized communities. For wage labor, we categorized each participant by whether or not they engaged in wage labor sometime during the past month. The diversity of wage labor conducted by the Orang Asli ranges from factory work, government service, work on plantations (oil palm and rubber), and resource extraction (timber and tin mining) (Gomes, 2004; Wallace et al., 2022).

Unlike previous studies of sleep among non-industrial societies, exposures of interest in this study did not include ecological factors such as daylight cycles and climatic conditions. This is because in Peninsular Malaysia, daylight hours and climate remain generally stable throughout the year. Located near the equator, the region experiences consistent photoperiods, with approximately 12 hours of daylight daily, fluctuating by less than 30 minutes annually. As a tropical environment, mean temperatures, humidity, and rainfall remain persistently high year-round. Thus, it is reasonable to assume that seasonality does not have a major impact on Orang Asli sleep.

Outcomes of interest included measures of sleep timing, quality, and quantity. For sleep timing, we examined the time of sleep onset and offset, as well as the Sleep Regularity Index (SRI), which measures the similarity of an individual’s sleep-wake patterns from one day to the next, with higher values indicating more regular sleep. For sleep quality, we focused on the number of awakenings lasting >5 minutes between sleep onset and offset, and waking after sleep onset (WASO), defined as the total time spent awake after sleep onset and before sleep offset. WASO is reflective of both the frequency and duration of nighttime awakenings, indicating sleep fragmentation. A lower WASO value indicates more consolidated, less disrupted sleep. For sleep quantity, we examined sleep duration, defined as the accumulated sustained nocturnal inactivity bouts within the sleep period (the time between sleep onset and offset).

### Statistical analyses

We used multilevel models to test for effects of exposures of interest on sleep outcomes (Table S1). Models for all outcomes used a Gaussian error distribution except for the number of awakenings (counts), for which we used a Poisson error distribution with log link function. Model structure was determined for each exposure variable using directed acyclic graphs (DAGs) to identify confounders and the appropriate set of adjustment variables for inclusion (Figure S1). In all models, observations represented person-nights and participant was included as a random intercept to adjust for repeat observations at the individual level.

Outcome variables were standardized (centered and divided by standard deviation) to facilitate comparison of effect sizes across models (Table S5). Model predictions were back-transformed and averaged across different levels of the predictor variables (or set at the average value of continuous predictors) to calculate estimated marginal means using the emmeans package in R (the Kenward-Rogers method was used to calculate degrees of freedom given the multilevel structure of models, and the Tukey method was used for post-hoc comparisons of variables with more than two levels).

All analyses were run using R version 4.5.0. All data and associated code is available on our GitHub (https://github.com/Kathleen-Reinhardt/sleep-circadian-lifestyle-malaysia.git).

## RESULTS

### Sleep outcomes vary between and within individuals

Together, we analyzed data from 5056 complete night recordings for 1036 individuals (N female= 669, N male= 367; average of 6.15 ± 1.67 nights per individual; Figure 2A and Table S2). These data broadly covered the adult life course, including participants aged 18-91 years old. Across all nights included in the dataset, individuals began sleeping on average at 11:53pm (+/-1.54 hours) and waking at 7:10am (+/-1.28 hours). On average, individuals experienced 17.7 awakenings (+/-6.03), corresponding to a wake after sleep onset time of 1.35 hours (+/-0.75 hours) and a total sleep duration of 5.92 hours (+/-1.43 hours). The average sleep regularity index was 55.20 (+/-18.61). As expected, most sleep variables were correlated, with the strongest positive relationships between awakenings and duration (Pearson correlation, R=0.605, p<10^−10^) as well as offset and duration (R=0.433, p<10^−10^; Figure 2B and Table S3). In other words, individuals who woke up more during the night, or who woke up later, slept for overall longer amounts of time. The strongest negative correlations were observed between onset and awakenings (R=-0.484, p<10^−10^) as well as onset and WASO (R=-0.363, p<10^−10^; Table S3). Individuals who went to sleep later thus woke up less during the night and had shorter wake after sleep onset times.

**Figure 2.**
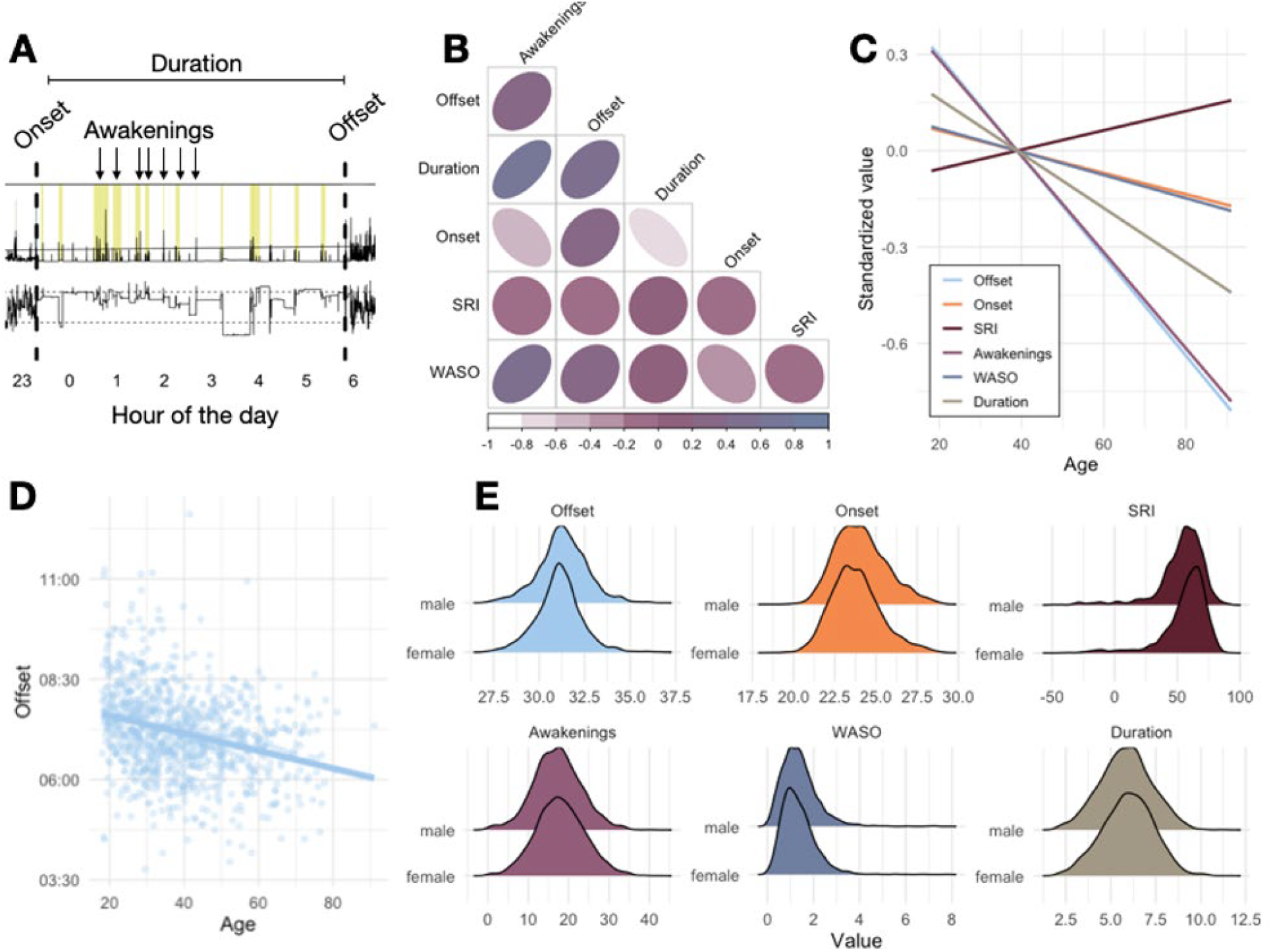
Overview of sleep variables and demographic effects on sleep timing, quality, and quantity. (A) Overview of select sleep variables assessed in this study and how they are extracted from an example accelerometry data file, specifically focusing on sleep offset (the time a person wakes up), sleep onset (the time a person goes to sleep), sleep duration (the total time asleep), and total number of awakenings. In downstream analyses, two composite variables, sleep regularity index (SRI) and wake after sleep onset (WASO), are also included. The upper plot represents acceleration and lower plot represents angle. (B) Correlation matrix showing pairwise relationships between the full set of sleep variables included. Ellipse shape and shading denote strength and direction of Pearson correlation coefficients. (C) Effects of age on standardized sleep outcomes (with each sleep variable scaled to mean 0 and unit variance). Lines represent best fit for each bivariate relationship. (D) Effect of age on sleep offset in unstandardized data, emphasizing that older individuals wake up earlier. Each point represents the average value for a given individual across all days of valid accelerometry data. (E) Sleep variable distributions broken down by sex. All plots use unstandardized values.

For all sleep outcomes, a substantial and statistically significant portion of the variance was attributable to individual differences (ANOVA, all p<10^−10^; Table S4). For example, 35% of the total variance in sleep regularity index was explained by differences between individuals, while the remaining 65% reflected within-individual variation across nights. SRI had the greatest within-individual variance and sleep offset had the lowest levels of within-individual variation, with an estimated 56% and 44% of the variance for these variables attributed to differences between versus within individuals, respectively.

### Sleep outcomes are structured by age and sex

Consistent with previous studies, we observed significant differences in sleep outcomes between sexes and across the life course. First, age impacts sleep timing and duration, with effects on onset, offset, and duration. Specifically, older individuals went to sleep earlier (β= -0.1, p<0.001), woke up earlier (β= -0.27, p<0.001), and overall slept for shorter amounts of time relative to younger individuals (β= -0.11, p<0.001; Figure 2C-D; Table S5). Age also impacted sleep quality: older individuals had shorter wake after sleep onset times (β= -0.05, p=0.046), higher sleep regularity indexes (β= 0.08, p<0.001), and fewer awakenings relative to younger individuals (incidence rate ratio=0.94, p<0.001).

Sex structured almost all of our analyzed sleep outcomes, except WASO. Males had later sleep onsets (β= 0.27, p<0.001) and offsets (β= 0.22, p<0.001), but shorter overall sleep durations (β= -0.12, p=0.009) relative to females (Figure 2E; Table S5). Males also experienced fewer awakenings relative to females (incidence rate ratio=0.96, p=0.017), but nonetheless exhibited lower sleep regularity indexes (β= -0.21, p<0.001).

### Electricity, but not housing type, technology, and wage labor, had consistent effects on sleep outcomes

After adjusting for age and sex, the availability of electricity exhibited major effects on the different sleep outcomes examined (Figs. 3,4). Electricity access had significant effects on the timing, quality, and quantity of sleep. For example, compared to those with no electricity, individuals with residential electricity from power lines went to sleep ∼28 minutes later (β= -0.31, SE= 0.07, *p* < 0.001), exhibited less sleep regularity (β= -0.24, SE= 0.06, *p* < 0.001), had 1.9 fewer awakenings per night (β= -0.11, SE= 0.02, *p* < 0.001) and spent 6.5 fewer minutes of waking time after sleep onset (β= -0.16, SE= 0.06, *p* = 0.01), and slept ∼23 fewer minutes per night (β= -0.26, SE= 0.06, *p* < 0.001) on average (note that reported effect sizes are standardized; Table S5). Access to electricity from generators had similar effects on sleep quality, and access to solar power had similar effects on sleep duration (Fig. 3). Electricity was thus associated with less sleep overall, but of generally higher quality (fewer awakenings and time spent awake after sleep onset).

**Figure 3.**
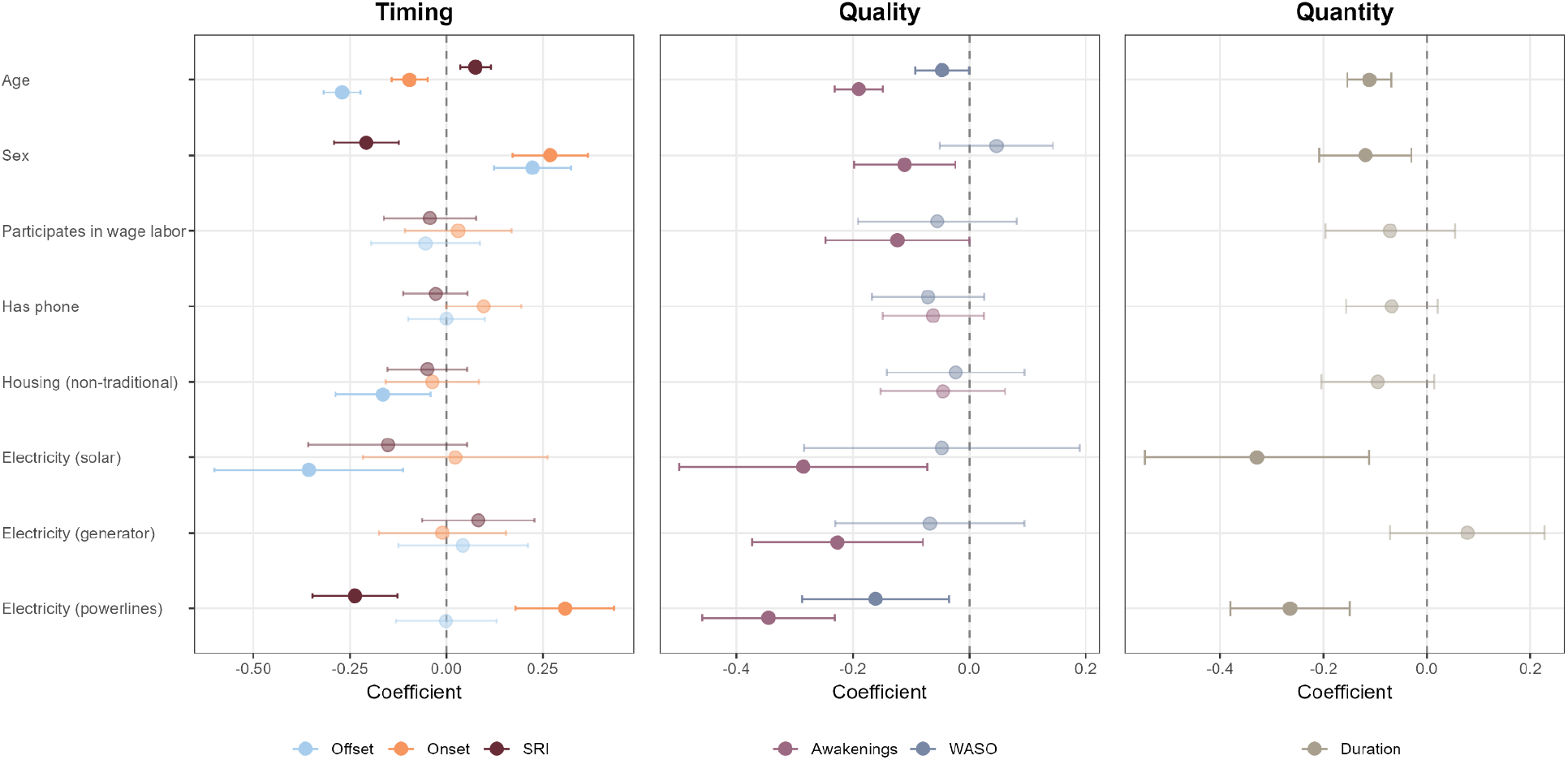
Comparison of effect size estimates for different variables affecting sleep outcomes. Points and error bars represent coefficients ± 95% CIs from hierarchical linear models for each sleep outcome, including all of the predictor variables along the y-axis. Response variables were standardized to facilitate comparability. Solid and transparent points represent significant (P < 0.05) and non-significant effects, respectively.

**Figure 4.**
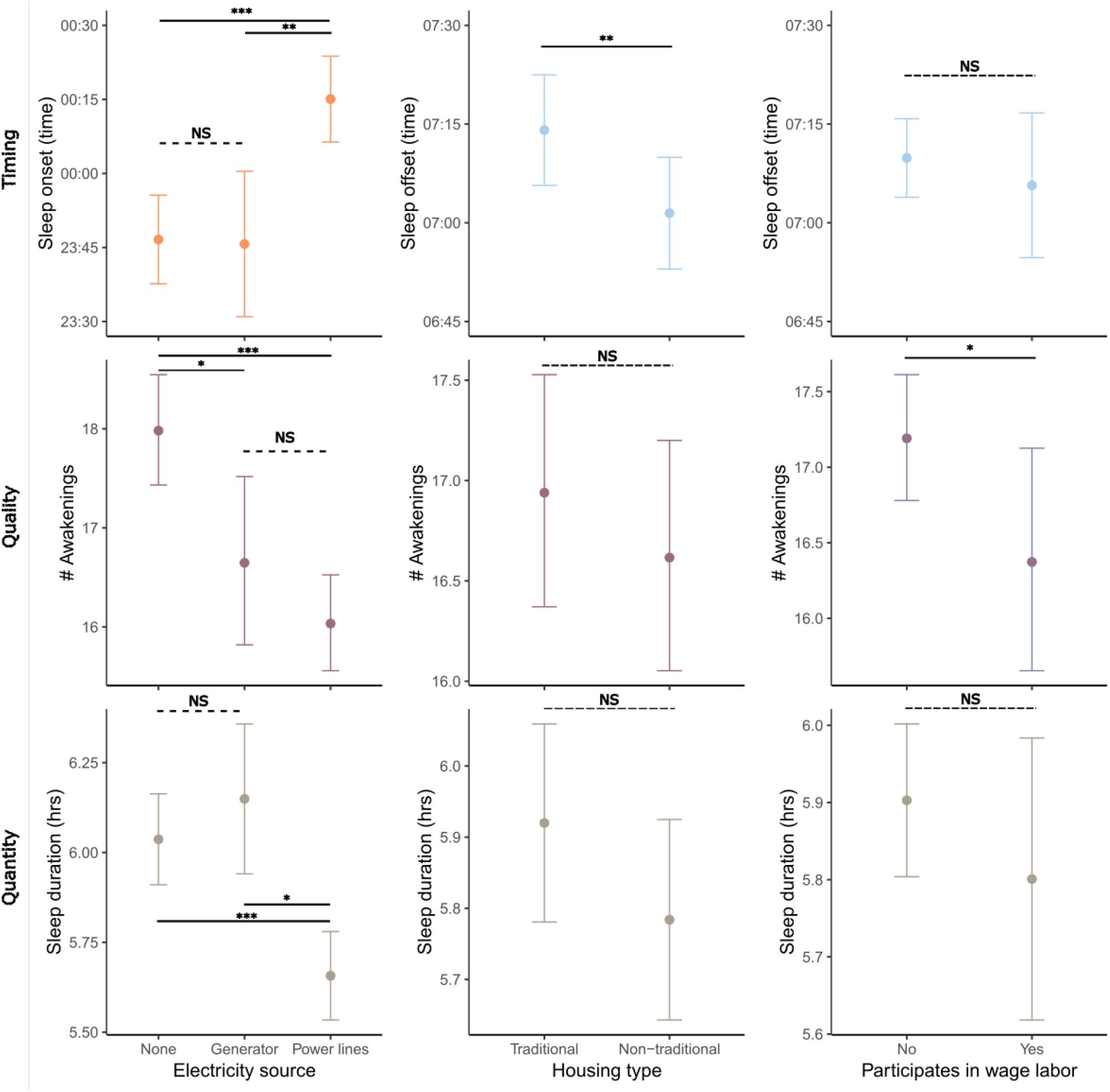
Estimated marginal mean differences in sleep outcomes as a function of predictors exhibiting effects of the greatest magnitude. Rows constitute different categories of sleep outcomes, columns show individual predictors. Points and error bars represent predicted marginal mean response ± 95% CIs on a natural scale (back-transformed when necessary). Significance from hierarchical model output is denoted as: *** = P < 0.001, ** = P < 0.01, * = P < 0.05, NS = not significant.

We observed no significant effects of owning a smartphone or different types of housing materials (concrete/wood vs. traditional bamboo thatch) on any aspect of sleep (Fig. 3, Table S5). Those participating in wage labor did exhibit slightly reduced nightly awakenings, but the result was small with high uncertainty (β= -0.05, SE= 0.02, *p* = 0.03), and this variable impacted neither sleep timing nor duration.

## DISCUSSION

Using a cross-sectional sample that comprises the largest study yet conducted on sleep among non-industrialized societies (n= 1036), we sought to examine how aspects of market integration (including modern technology, built environment, and labor practices) shape sleep timing, quality, and quantity among the Orang Asli of Peninsular Malaysia, an Indigenous population undergoing significant lifestyle changes. We found that constant electricity had robust effects on sleep timing, quality, and quantity while permanent housing and wage labor participation showed weaker associations. These findings extend the comparative literature on sleep in non-industrialized societies.

### Electricity source impacts sleep timing, quality, and quantity

Consistent with cross-cultural research, we found that access to electricity via powerlines delayed sleep onset and increased timing irregularity among the Orang Asli (de la Iglesia et al. 2015; Louzada et al. 2024; Pilz et al. 2018; Smit et al. 2019). This aligns with broader evidence that electricity weakens circadian alignment to natural zeitgebers and expands social activity at night. In contrast, access to solar electricity and generators (which are limited in power run-time) had no impact on sleep timing, paralleling findings that both the type and autonomy of electricity access impact circadian rhythms as much as the mere presence of light (Duffy & Wright, 2005; Peirson et al. 2018; Roenneberg & Merrow, 2007). Although smartphones have proliferated among the Orang Asli over the past ten years, smartphone ownership was not linked with delayed sleep phasing, offering contrasting evidence from clinical and industrial contexts (Chang et al 2015; Delorme et al 2022). This suggests that habitual use and the timing of smartphone exposure, rather than ownership per se, are likely more consequential.

Somewhat paradoxically, powerline access resulted in fewer awakenings and lower WASO, suggesting significantly better sleep quality among more urbanized Orang Asli. However, higher sleep quality occurred alongside shorter sleep duration. This was due to later sleep onset yet no corresponding shift in sleep offset. One explanation for this pattern is that later sleep onset increases homeostatic sleep pressure, leading to more consolidated and less fragmented sleep. This process has been documented in the literature (Vyazovskiy & Tobler 2012, Vyazovskiy et al. 2017), but not yet in any small-scale society. This finding aligns with cross-cultural evidence that electricity access narrows the sleep period (the time between sleep onset and sleep offset) without always compromising the quality of sleep (de la Iglesia et al. 2015; Pilz et al. 2018; Smit et al. 2019; Samson, 2021).

### The built environment minorly impacts sleep timing but not quality or quantity

Despite their increasing presence in Orang Asli villages, non-traditional concrete or wood houses are almost universally disliked by the Orang Asli, as they often lack functional sanitation, are prone to degradation from stagnant water, are often overcrowded with people, and have limited airflow, making them very hot (Latif & Essah 2024, Abdullah et al 2016, Hamdan et al 2007). In contrast, traditional houses are constructed from natural materials and are raised off the ground, resulting in adequate ventilation. In other words, traditional houses are well-suited to the hot tropical environment (Majid & Rosali 2025). As such, we expected non-traditional, government-constructed houses to blunt natural light and temperature cues, delaying sleep timing of both onset and offset (Mistlberger & Skene, 2005; Wright, 2005). However, individual waking time shifted earlier by only 10 minutes, a biologically negligible effect. Contrary to previous research linking more permanent housing to higher sleep efficiency (McKinnon et al., 2023; Nunn & Samson, 2016; Samson et al., 2017,Samson et al., 2025; Yetish et al 2015), we found no evidence that housing type influenced sleep quality or quantity. Overall, despite their lack of comfort during the daytime, non-traditional housing does not appear to significantly impact Orang Asli sleep.

### Wage labor modestly impacts sleep quality but not timing or quantity

While labor practices may impact sleep patterns due to trade-offs between sleep and time spent working (Prall et al., 2018), among the Orang Asli, participation in wage labor had no effect on sleep timing or quantity. It was, however, associated with a modest (5%) reduction in nighttime awakenings. While the effect size is small, this aligns with literature that links high amounts of daytime physical activity (of low and moderate intensity) with higher sleep quality (Wang & Boros, 2021; Zhao et al., 2023) including among other non-industrialized societies (i.e. Li et al., 2025). Our anecdotal observations suggest that wage labor as practiced among the Orang Asli is more time-structured, repetitive and physically demanding as compared to traditional subsistence practices. Despite shifts to more time-structured wage labor, no impact on regularity on the timing of sleep was evident in our data. Yet, given the physically intense nature of wage labor among the Orang Asli, it is possible that labor changes can subtly affect sleep quality. Taken together, these data suggest that while wage labor is not a primary driver of sleep timing or quantity among the Orang Asli, it may exert secondary effects on quality through sleep consolidation and homeostatic sleep pressure.

### Age and sex are strong biological and ecological determinants of Orang Asli sleep

Age and sex emerged as strong and consistent predictors of sleep and circadian rhythm variation. Older individuals exhibited earlier, and more regular and shorter sleep, consistent with well-documented circadian phase advances (earlier to bed, earlier to rise) that are associated with aging (Espiritu, 2008; Li et al., 2022; Mander et al., 2006; Ohayon et al., 2004; Vitiello, 2006). However, in contrast to this literature, we did not observe an increase in fragmentation that is typically reported among aging adults in industrialized settings (Samson & McKinnon, 2025). Rather, we observed a decrease in both nighttime awakenings and WASO with age. This suggests that while age-related decline in sleep duration may indeed be a universal phenomenon, in non-industrial populations it could reflect adaptive homeostasis rather than the pathological increase in nighttime awakenings reported in industrial settings. As such, while the functional consequences of reduced sleep duration among older individuals remain unclear, our findings add to a growing body of evidence suggesting that sleep needs may shift with age (Ohayon et al. 2004).

We also observed sex differences in sleep behavior. Compared to women, men displayed later sleep onset and offset, more irregular timing, overall shorter duration, and fewer nighttime awakenings. Among the Orang Asli, this likely reflects gendered labor patterns, where men are more likely to engage in physically demanding tasks such as hunting and wage labor (Endicott & Endicott 2007), leading to increased sleep pressure and likely contribute to later, more irregular sleep timing of a shorter duration. These patterns are mixed in comparison to other non-industrialized societies where sex-specific divisions of labor appear to structure daily activities, matching other hunter-gatherer and horticulture subsistence societies (Kilius et al., 2021; Yetish et al., 2018), and contrasting with pastoralist societies in which men experience lower sleep quality (Prall et al., 2018). These comparisons underscore how diverse ecologies and culturally mediated divisions of labor shape sleep beyond biological sex effects.

### Environmental impacts on Orang Asli sleep

We observed very late sleep onset among the Orang Asli, a few hours later than has been documented in other non-industrial societies (de la Iglesia et al., 2018; Prall et al., 2018; Yetish et al., 2015). Although we did not measure environmental variables directly in our study, the tropical environment occupied by Orang Asli in Peninsular Malaysia may help to explain some of the sleep patterns we observed (Fig. 1D). Daytime temperatures are extremely high (mid 20s-low 30s C), which could discourage activity, whereas nighttime temperatures drop by up to 11 degrees Celsius (depending on elevation). Orang Asli may go to bed late simply because evening is the only time to comfortably move around and socialize. Also, Orang Asli rise to begin their day when temperatures reach their coolest at around 7AM. As temperature can impact circadian organization via effects on sleep-wake and activity patterns (Mistlberger & Skene, 2005), our data show a tight association between sleep timing and social rhythms in the tropical climate of Malaysia.

### Cross-cultural comparisons of sleep variation

Cross-culturally, sleep efficiency appears to be higher in industrial populations, suggesting better sleep quality compared to non-industrial populations (Samson & McKinnon, 2025). Compared to clinical studies and epidemiological studies on both industrial and other non-industrial societies, Orang Asli showed slightly higher WASO (median: 1.25 hrs) and more awakenings (median: 18 per night) (Dashti et al. 2019; Ohayon et al. 2004). It is important to note that the number of awakenings estimated by actigraphy likely overestimates true physiological or perceived awakenings, as brief movement-induced wake bouts are counted as separate events (Ancoli-Israel et al 2015). However, as we applied the same scoring algorithm and thresholds consistently across all participants, this acts as an internal control, making the relative differences informative. What is particularly notable among the Orang Asli is the direction of these associations. As individuals adopt more urbanized lifestyles, they exhibit lower WASO and fewer prolonged awakenings, suggesting improved sleep consolidation. Furthermore, the observed decrease in both nighttime awakenings and WASO with age diverges from widely reported patterns of increased sleep fragmentation and decreased sleep efficiency in older adults (Espiritu, 2008; Mander et al. 2017).

The Orang Asli slept an average of 6 hours per night; a duration on the lower end of global distributions but consistent with recent findings that non-industrial populations often sleep less than industrial populations (Samson & McKinnon, 2025). This relatively short sleep quantity occurs along with elevated sleep quality, supporting the argument that short sleep is not inherently pathological when embedded in ecological and cultural context.

### Links between sleep and health

Dysregulation of the circadian clock is thought to play a causal role in numerous pathologies (Roennenberg & Merrow, 2016). However, the association between health outcomes and sleep quantity remains unclear. One limitation of the present study is that we did not measure the self-reported sleep quality of participants, thus, we do not know the potential performance and downstream health costs of poorer sleep, or whether participants in our study evinced sleep disorders. Ou et al (2025) found that sleep duration and health were related in a quadratic fashion across several countries, with individuals whose sleep duration was closer to their country’s perceived ideal reporting better health. Ou et al (2025) conclude that a good night of sleep is not only related to physiological need but also shaped by cultural standards. Evidence from the Orang Asli supports this idea. The average 6 hours of sleep displayed among the Orang Asli lies near the optimum value for minimizing mortality risk (Shen et al 2016), and while less sleep is often associated with increased risk for cardiovascular disease (Wu et al., 2025), the Orang Asli generally exhibit good cardiometabolic health (Watowich et al., 2024, Watowich et al., 2025). While preliminary, these associations offer support for Ou et al’s (2025) supposition and challenge the idea that deviation from 7-8 hour sleep is universally harmful.

## Conclusion

This comprehensive study illustrates how market integration impacts sleep and circadian rhythms, offering insight into the proximate mechanisms by which sleep disorders may arise. Sleep timing, quality, and quantity of the Orang Asli are most impacted by access to permanent electricity sources and individual-level factors such as age and sex, and less so by novel labor practices. These results point to the need for an interdisciplinary approach that considers how biological, ecological, and social factors interact within cultural frameworks to shape global sleep disorders.

## Acknowledgements

We thank the Orang Asli communities for their participation in and support for our research. We are grateful to all past and present members of the Orang Asli Health and Lifeways Project and the Malaysian Red Crescent Society for providing research support for data collection. We are also grateful to the Malaysian Economic Planning Unit and the Department of Orang Asli Development (JAKOA) for their support.

## Ethics

Procedures for this study have been reviewed and approved by the Medical Review and Ethics Committee of the Malaysian Ministry of Health (protocol ID: NMRR-20-2214-55565), the Malaysian Ministry of Economy (permit ID: EPU 40/200/19/3911), the Malaysian Department the of Orang Asli Development (permit ID: JAKOA.PP.30.052 JLD 21 (98)) and the Institutional Review Boards of the University of New Mexico (protocol ID: 14420), Vanderbilt University (protocol ID: 212175), and the University of Calgary (REB21-0432). Throughout the project, we have followed established principles for ethical biomedical research among Indigenous communities, including fostering collaboration, building cultural competency, being transparent about research practices, supporting capacity building, and disseminating research findings (Claw et al 2018). Informed consent was obtained from all participants at both the community and individual level for this study. The study procedures and rationale were described to participants in the language of choice (Malay or the local Orang Asli language), supplemented by translated printouts of information for those wishing to read a physical copy.

