## Supplementary Material for "Recent lifestyle change impacts sleep and circadian rhythms among the Indigenous peoples of Peninsular Malaysia"

Table S1. Reported access to powerlines, living in non-traditional housing, and participating in wage labor across study locations.

| | Number of individuals asked | Number that answered yes | Proportion that answered yes |
|---|---|---|---|
| Has powerlines | 954 | 430 | 0.4507337526 |
| Lives in non-traditional housing | 944 | 510 | 0.5402542373 |
| Works wage labor | 943 | 127 | 0.1346765642 |

Table S2. Summary of participant sleep characteristics. Values are presented as means (and standard error) for the complete data sample, separately by sex. Abbreviations: WASO, waking after sleep onset; SRI, sleep regularity index. Hours are presented as decimals and time is presented as HH:MM.

| | **males** | **females** |
|---|---|---|
| N total participants (nights) | 367 (1808 nights) | 669 (3248 nights) |
| Avg. nights per individual | 6.22 ± 1.74 | 6.12 ± 1.62 |
| Age range (years) | 18-81 | 18-91 |
| Sleep onset (hh:mm) | 00:06 ± 1:34 | 23:47 ± 1:30 |
| Sleep offset (hh:mm) | 07:19 ± 0:31 | 07:06 ± 0:43 |
| Sleep period (hours) | 7.22 ± 1.67 | 7.32 ± 1.68 |
| Sleep duration (hours) | 5.83 ± 1.44 | 5.98 ± 1.43 |
| WASO (Waking After Sleep Onset) | 1.39 ± 0.78 | 1.34 ± 0.73 |
| SRI (Sleep Regularity Index) | 53.5 ± 18.4 | 56.2 ± 18.2 |

Table S3. Pearson correlation coefficients (top) and p-values (bottom) for pairwise comparisons between sleep variables.

| | Awakenings | Offset | Duration | Onset | SRI | WASO |
|---|---|---|---|---|---|---|
| Awakenings | 1 | 0.3407086 | 0.60532366 | -0.48429287 | -0.009596321 | 0.42320798 |
| Offset | 0.340708597 | 1 | 0.43274979 | 0.31080785 | -0.086945 | 0.2441266 |
| Duration | 0.605323664 | 0.4327498 | 1 | -0.60911228 | 0.071372209 | 0.08520901 |
| Onset | -0.48429287 | 0.3108078 | -0.60911228 | 1 | -0.078257361 | -0.36326084 |
| SRI | -0.009596321 | -0.086945 | 0.07137221 | -0.07825736 | 1 | -0.12114171 |
| WASO | 0.42320798 | 0.2441266 | 0.08520901 | -0.36326084 | -0.121141713 | 1 |

| | Awakenings | Offset | Duration | Onset | SRI | WASO |
|---|---|---|---|---|---|---|
| Awakenings | 0.00E+00 | 3.54E-136 | 0.00E+00 | 1.49E-292 | 5.55E-01 | 1.39E-216 |
| Offset | 3.54E-136 | 0.00E+00 | 1.90E-227 | 1.81E-112 | 8.71E-08 | 8.86E-69 |
| Duration | 0.00E+00 | 1.90E-227 | 0.00E+00 | 0.00E+00 | 1.13E-05 | 1.58E-09 |
| Onset | 1.49E-292 | 1.81E-112 | 0.00E+00 | 0.00E+00 | 1.47E-06 | 6.97E-156 |
| SRI | 5.55E-01 | 8.71E-08 | 1.13E-05 | 1.47E-06 | 0.00E+00 | 8.01E-14 |
| WASO | 1.39E-216 | 8.86E-69 | 1.58E-09 | 6.97E-156 | 8.01E-14 | 0.00E+00 |

Table S4. ANOVA model estimates of the proportion of variance in sleep outcomes explained by between- vs. within-individual differences.

| Outcome | Sum of squares (between) | Sum of squares (within) | PVE between | PVE within | F value |
|---|---|---|---|---|---|
| Awakenings | 84887.338 | 97033.89 | 0.4666159 | 0.5333841 | 3.557785 |
| Offset | 4559.073 | 3608.196 | 0.5582127 | 0.4417873 | 5.138622 |
| Duration | 4495.209 | 5673.044 | 0.4420827 | 0.5579173 | 3.222508 |
| Onset | 5966.305 | 5860.907 | 0.5044557 | 0.4955443 | 4.140005 |
| SRI | 457753.168 | 849303.992 | 0.3502166 | 0.6497834 | 1.675571 |
| WASO | 1276.913 | 1529.492 | 0.4549996 | 0.5450004 | 3.395271 |

Table S5. Standardized linear mixed model estimates for each sleep outcome, showing fixed effects of predictors (top) and model variance components (bottom).

| | Sleep offset | | | | Sleep onset | | | | SRI | | | | # Awakenings (Poisson GLMM) | | | | WASO | | | | Duration | | | |
|---|---|---|---|---|---|---|---|---|---|---|---|---|---|---|---|---|---|---|---|---|---|---|---|---|
| *Predictors* | *Estimates* | *std. Error* | *CI* | *p* | *Estimates* | *std. Error* | *CI* | *p* | *Estimates* | *std. Error* | *CI* | *p* | *Incidence Rate Ratios* | *std. Error* | *CI* | *p* | *Estimates* | *std. Error* | *CI* | *p* | *Estimates* | *std. Error* | *CI* | *p* |
| Intercept | 0.03 | 0.05 | -0.07 – 0.13 | 0.553 | -0.26 | 0.05 | -0.36 – -0.17 | <0.001 | 0.24 | 0.04 | 0.16 – 0.32 | <0.001 | 19.12 | 0.3 | 18.55 – 19.71 | <0.001 | 0.13 | 0.05 | 0.04 – 0.22 | 0.007 | 0.25 | 0.04 | 0.17 – 0.34 | <0.001 |
| Age | -0.27 | 0.02 | -0.32 – -0.22 | <0.001 | -0.1 | 0.02 | -0.14 – -0.05 | <0.001 | 0.08 | 0.02 | 0.04 – 0.12 | <0.001 | 0.94 | 0.01 | 0.92 – 0.95 | <0.001 | -0.05 | 0.02 | -0.09 – -0.00 | 0.046 | -0.11 | 0.02 | -0.15 – -0.07 | <0.001 |
| Sex (1=male; 0=female) | 0.22 | 0.05 | 0.12 – 0.32 | <0.001 | 0.27 | 0.05 | 0.17 – 0.37 | <0.001 | -0.21 | 0.04 | -0.29 – -0.12 | <0.001 | 0.96 | 0.02 | 0.93 – 0.99 | 0.017 | 0.05 | 0.05 | -0.05 – 0.14 | 0.343 | -0.12 | 0.05 | -0.21 – -0.03 | 0.009 |
| Participates in wage labor | -0.05 | 0.07 | -0.19 – 0.09 | 0.452 | 0.03 | 0.07 | -0.11 – 0.17 | 0.66 | -0.04 | 0.06 | -0.16 – 0.08 | 0.489 | 0.95 | 0.02 | 0.91 – 1.00 | 0.031 | -0.05 | 0.07 | -0.19 – 0.08 | 0.43 | -0.07 | 0.06 | -0.20 – 0.05 | 0.267 |
| Has phone | 0 | 0.05 | -0.10 – 0.10 | 0.985 | 0.1 | 0.05 | -0.00 – 0.19 | 0.052 | -0.03 | 0.04 | -0.11 – 0.06 | 0.519 | 0.98 | 0.02 | 0.95 – 1.01 | 0.294 | -0.07 | 0.05 | -0.17 – 0.03 | 0.149 | -0.07 | 0.05 | -0.16 – 0.02 | 0.133 |
| Housing (non-traditional) | -0.16 | 0.06 | -0.29 – -0.04 | 0.009 | -0.04 | 0.06 | -0.16 – 0.08 | 0.557 | -0.05 | 0.05 | -0.15 – 0.06 | 0.359 | 0.98 | 0.02 | 0.94 – 1.02 | 0.323 | -0.02 | 0.06 | -0.14 – 0.10 | 0.7 | -0.09 | 0.06 | -0.20 – 0.01 | 0.089 |
| Electricity (solar) | -0.36 | 0.13 | -0.60 – -0.11 | 0.004 | 0.02 | 0.12 | -0.22 – 0.26 | 0.849 | -0.15 | 0.11 | -0.36 – 0.06 | 0.151 | 0.92 | 0.04 | 0.85 – 0.99 | 0.028 | -0.05 | 0.12 | -0.28 – 0.19 | 0.696 | -0.33 | 0.11 | -0.55 – -0.11 | 0.003 |
| Electricity (generator) | 0.04 | 0.09 | -0.12 – 0.21 | 0.614 | -0.01 | 0.08 | -0.17 – 0.15 | 0.906 | 0.08 | 0.07 | -0.06 – 0.23 | 0.263 | 0.93 | 0.02 | 0.88 – 0.98 | 0.004 | -0.07 | 0.08 | -0.23 – 0.10 | 0.416 | 0.08 | 0.08 | -0.07 – 0.23 | 0.305 |
| Electricity (powerlines) | 0 | 0.07 | -0.13 – 0.13 | 0.993 | 0.31 | 0.07 | 0.18 – 0.44 | <0.001 | -0.24 | 0.06 | -0.35 – -0.13 | <0.001 | 0.89 | 0.02 | 0.86 – 0.93 | <0.001 | -0.16 | 0.06 | -0.29 – -0.03 | 0.012 | -0.26 | 0.06 | -0.38 – -0.15 | <0.001 |

| Random Effects | Sleep offset | Sleep onset | SRI | # Awakenings (Poisson GLMM) | WASO | Duration |
|---|---|---|---|---|---|---|
| $\sigma^2$ | 0.55 | 0.63 | 0.74 | 0.06 | 0.64 | 0.7 |
| $\tau_{00}$ | 0.4 | 0.36 | 0.15 | 0.04 | 0.34 | 0.26 |
| ICC | 0.42 | 0.36 | 0.16 | 0.41 | 0.35 | 0.27 |
| N | 979 | 979 | 916 | 979 | 979 | 979 |
| Observations | 4755 | 4755 | 3580 | 4755 | 4755 | 4755 |
| Marginal $R^2$ / Conditional $R^2$ | 0.088 / 0.473 | 0.041 / 0.390 | 0.033 / 0.192 | 0.099 / 0.465 | 0.014 / 0.359 | 0.052 / 0.306 |
