## Supplementary material for "Recent lifestyle change impacts sleep and circadian rhythms among the Indigenous peoples of Peninsular Malaysia": SI Appendices

**Table 1.** Hypotheses and adjusted sets derived from DAGs to structure each testable model.

| **Hypothesis: Exposure** | **Minimal adjusted set (for confounders)** | **Refined model (with interactions)** |
|---|---|---|
| **H0:** Null Hypothesis | None (random intercept only) "Individual differences" | Sleep output ~ (1\|rid) + (1\|location) |
| **H1:** Age | None https://dagitty.net/dags.html?id=nxmpmJez | Sleep output ~ age +(1\|rid) + (1\|location) |
| **H2:** Sex | None https://dagitty.net/dags.html?id=AhSqaJV9 | Sleep output ~ sex +(1\|rid) + (1\|location) |
| **H3:** Electricity | Housing + Wage labor https://dagitty.net/dags.html?id=9cPTgLhK | Sleep output ~ electricity_source + house_type + wage_past_month + (1\|rid) + (1\|location) -plausible interactions: electricity*house type, electricity*wage labor |
| **H4:** Smart phone | Age + Electricity + Wage labor https://dagitty.net/dags.html?id=cAdpE4RZ | Sleep output ~ smart_phone + age + electricity_source + wage_past_month + (1\|rid) + (1\|location) -plausible interactions: |
| **H5:** House type | Wage labor + Age https://dagitty.net/dags.html?id=i3eMgKWr | Sleep output ~ house_type + age + wage_past_month + (1\|rid) + (1\|location) -plausible interactions: |
| **H6:** Wage labor | Age + Sex https://dagitty.net/dags.html?id=EYiqSYU9 | Sleep output ~ wage_past_month + age + sex + (1\|rid) + (1\|location) |

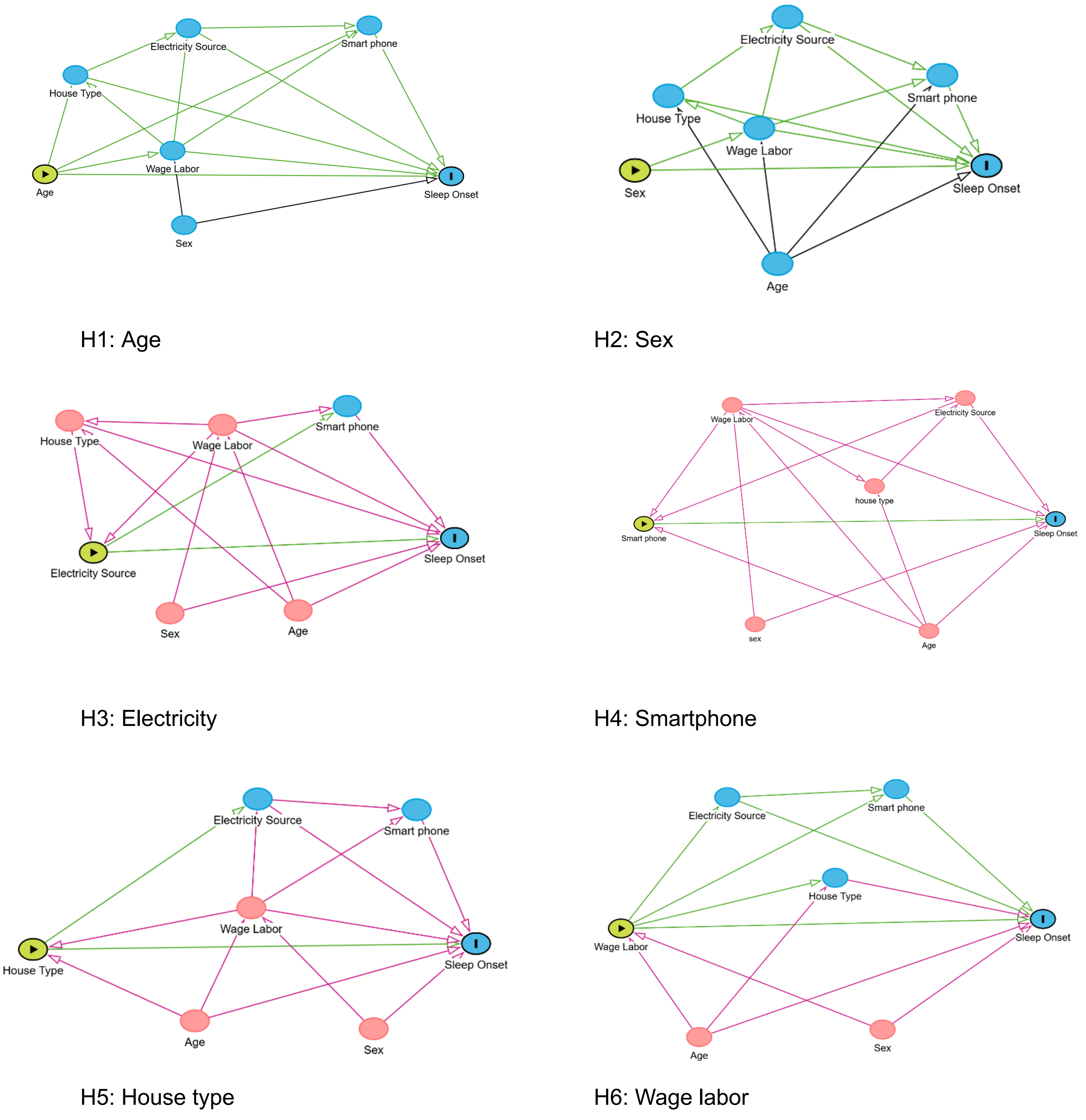


**S1**. DAGs for each lifestyle variable.
